# NLRP3 inflammasome inhibition rescues Hutchinson-Gilford Progeria cellular phenotype and extend longevity of an animal model

**DOI:** 10.1101/2020.09.09.288290

**Authors:** Elísabet Alcocer-Gómez, Beatriz Castejón-Vega, Jéssica Nuñez-Vasco, Débora Lendines-Cordero, José M. Navarro-Pando, Mario D. Cordero

## Abstract

Inflammation is a hallmark of aging and accelerated aging syndromes. In this context, inflammation has been associated to the pathophysiology of Hutchinson–Gilford progeria syndrome (HGPS). In this study, we report that progeroid skin fibroblasts and animal models present an hyperactivation of the NLRP3-inflammasome complex. High expression of NLRP3 and caspase 1 was also observed in skin fibroblasts from HGPS associated to the nuclei morphology. Lymphoblast from HGPS also showed increased basal levels of NLRP3 and caspase 1 independent to the induction from metabolic factors. Consistent with these results, Zmpste24^−/−^ showed high expression of Nlrp3 and caspase 1 in heart, liver and kidney and reduced levels of Nlrc3, however these changes were not observed in other inflammasomes. We also show that pharmacological inhibition of NLRP3 using a direct NLRP3 inhibitor, MCC950, improved cellular phenotype, significantly extends the lifespan of these progeroid animals and reduced inflammasome-dependent inflammation. These findings suggest the NLRP3-inflammasome comples as a therapeutic approach for patients with HGPS.

---

Ageing involves a progressive impairment of physiological homeostasis, situation that is reflected in the cell tissues, and at organismal level (1). In this context, inflammation is highly associated to the aging process and age-related diseases and the inflamm-ageing, a low-grade sterile chronic inflammation, has been described as a progressive event during biological ageing with accumulation of pro-inflammatory mediators (2). In the last years, the role of NLRP3-inflammasome has been studied in many age-related diseases.

The NLRP3-inflammasome is one of the most well-studied inflammasomes in humans and mice (3). It is a multiprotein complex comprising NLRP3 itself as an intracellular sensor, the adapter protein ASC and pro-caspase-1. The NLRP3 inflammasome is activated by a range of danger and stress signals (3), some of which rise during aging (4). The Nlrp3 ablation in mice has been shown to improves lifespan and health by attenuating multiple age-related degenerative changes such as cardiac aging, insulin sensitivity, bone loss, and ovarian aging (5–7). According to this, the role of NLRP3-inflammasome has been assessed in normal aging yet remain unexplored in genetic models of accelerated aging. Hutchinson-Gilford progeria syndrome (HGPS) is a rare premature aging condition in which a point mutation in the LMNA gene (c.1824C > T; GGC > GCT; p.G608G) (8) causes the accumulation at the nuclear envelope of an aberrant precursor of lamin A, named progerin, which disrupts the nuclear membrane architecture and causes multiple cellular alterations, including abnormal gene transcription and signal transduction. The clinical phenotype is characterized by delayed loss of primary teeth, alopecia, osteoporosis, abnormal skin pigmentation, accelerated cardiovascular disease, growth impairment, lipodystrophy, dermal and bone abnormalities, and metabolic alterations (9). Inflammation has been associated to the pathophysiology of progeroid syndromes. Nuclear factor κB (NF-κB)-mediated secretion of high levels of proinflammatory cytokines have been shown in two different mouse models (10).

However, the genetic and pharmacological inhibition of NF-κB signaling prevented age-associated features in these animal models, and extended their longevity (10). Interestingly, NF-κB is a central mediator of the priming signal of NLRP3-inflammasome complex (11). In the present work, we report that human skin fibroblasts and lymphocytes from patients with HGPS and Zmpste24^−/−^ mice, an appropriate murine model of HGPS, demonstrate a clear activation of the NLRP3-inflammasome complex. In addition, we show that this alteration is detrimental, given that skin fibroblasts from patients treated with MCC950, a specific inhibitor of NLRP3, show improved phenotype and Zmpste24^−/−^ mice treated with MCC950 show a clear amelioration of progeroid features and inflammation and extended longevity compared with untreated Zmpste24^−/−^ mice.

## Material and methods

### Reagents

Trypsin was purchased from Sigma Chemical Co., (St. Louis, Missouri). Anti-actin monoclonal antibody from Calbiochem-Merck Chemicals Ltd. (Nottingham, UK). Lamin A/C, NLRP3, NLRC3 and caspase 1 were obtained from Cell Signaling Technology. MAP-LC3, NLRP4, Nalp1 and Nalp10 were obtained from Santa Cruz Biotechnology. NLRP3 inhibitors MCC950 and 16673-34-0 were obtained from Sigma-Aldrich (Saint Louis, USA). A cocktail of protease inhibitors (complete cocktail) was purchased from Boehringer Mannheim (Indianapolis, IN). Grace’s insect medium was purchased from Gibco. The Immun Star HRP substrate kit was from Bio-Rad Laboratories Inc. (Hercules, CA).

### Fibroblast culture

All fibroblasts from patients with HGPS were obtained from The Progeria Research Foundation Cell and Tissue Bank (http://www.progeriaresearch.org). The following fibroblasts were used: HGADFN367 (3-year-old male) and HGADFN155 (1.2-year-old female). Control fibroblasts were obtained from the Coriell Institute for Medical Research (Camden, NJ, USA). Fibroblasts were cultured in high glucose DMEM (Dulbecco’s modified media) (Gibco, Invitrogen, Eugene, OR, USA) supplemented with 15% fetal bovine serum (FBS) (Gibco, Invitrogen, Eugene, OR, USA), 1% GlutaMAX (ThermoFisher) and antibiotics (Sigma Chemical Co., St. Louis, MO, USA). Cells were incubated at 37°C in a 5% CO_2_ atmosphere. The medium was changed every two days to avoid changes in pH.

We also used the following lymphoblasts: HGALBV009 (5.1-year-old male) and HGALBV021 (father of the proband, 37-year-old female). Lymphoblasts were cultured in RPMI-1640 (Gibco, Invitrogen, Eugene, OR, USA) supplemented with 15% fetal bovine serum (FBS) (Gibco, Invitrogen, Eugene, OR, USA), and antibiotics (Sigma Chemical Co., St. Louis, MO, USA). Cells were incubated at 37°C in a 5% CO_2_ atmosphere

### Western Blotting

Whole cellular lysate from fibroblasts was prepared by gentle shaking with a buffer containing 0.9% NaCl, 20 mM Tris-ClH, pH 7.6, 0.1% Triton X-100, 1 mM phenylmethylsulfonylfluoride and 0.01% leupeptin. The protein content was determined by the Bradford method. Electrophoresis was carried out in a 10–15% acrylamide SDS/PAGE and proteins were transferred to Immobilon membranes (Amersham Pharmacia, Piscataway, NJ). Next, membranes were washed with PBS, blocked over night at 4°C and incubated with the respective primary antibody solution (1:1000). Membranes were then probed with their respective secondary antibody (1:2500). Immunolabeled proteins were detected by chemiluminescence method (Immun Star HRP substrate kit, Bio-Rad Laboratories Inc., Hercules, CA). Western blot images were quantified using ImageJ software.

### NLRP3 immunofluorescence

NLRP3 distribution in cytosol was assessed by immunofluorescence techniques using antibodies against NLRP3 and DAPI as a marker of the nuclei.

### Proliferation rate

Two hundred thousand fibroblasts were cultured with or without the MCC950 at two different concentrations (0.6 and 1.2mM) for 24, 48, and 120h. After discharging supernatant with dead cells, cells from three high-power fields were counted with an inverted microscope using a 40X objective.

### Animals

Animal studies were performed in accordance with European Union guidelines (2010/63/EU) and the corresponding Spanish regulations for the use of laboratory animals in chronic experiments (RD 53/2013 on the care of experimental animals). All experiments were approved by the local institutional animal care committee. For all experiments, only male mice were used. Mutant mice deficient in Zmpste24 metalloproteinase have been described previously (10). All groups had *ad libitum* access to their prescribed diet and water throughout the whole study. Body weight was monitored weekly. Animal rooms were maintained at 20–22°C with 30–70% relative humidity.

For all experiments with NLRP3 inhibitors, Zmpste24^+/+^ (wild type) and Zmpste24^−/−^ were maintained on a regular 12 h light/dark cycle at 20–22°C. Treatments were started at 1month of age after randomization into three groups (wild type vehicle, Zmpste24^−/−^ vehicle and Zmpste24^−/−^MCC950). These groups correspond to the following treatment: i) standard diet with i.p vehicle (saline) treatment (vehicle groups) from Teklad Global 14% Protein Rodent Maintenance Diet, Harlan Laboratories (carbohydrate:protein:fat ratio of 48:14:4 percent of kcal) and ii) standard diet with MCC950 treatment (MCC950 group). MCC950 was administered 20mg/kg daily by i.p. route. All groups had *ad libitum* access to their prescribed diet and water throughout the study. Individuals were monitored daily and weighed monthly but were otherwise left undisturbed until they died. Survival was assessed using male mice, and all animals were dead by the time of this report. Kaplan–Meier survival curves were constructed using known birth and death dates, and differences between groups were evaluated using the logrank test. A separate group of male mice were sacrificed at age 4 months to study (western blots).

### Statistical Analysis

Data in the figures is shown as mean ± SD. Data between different groups were analysed statistically by using ANOVA on Ranks with Sigma Plot and Sigma Stat statistical software (SPSS for Windows, 19, 2010, SPSS Inc. Chicago, IL, USA). For cell-culture studies, Student’s t test was used for data analyses. A value of p<0.05 was considered significant

## Results

### HGPS patients show NLRP3-inflammasome hyperactivation

To evaluate the role of the NLRP3-inflammasome in progeria syndrome, we examined the NLRP3-inflammasome complex expression in HGPS fibroblasts and age- and passage-matched controls. As shown in Fig. 1A, HGPS demonstrates a significant increment in NLRP3 and caspase 1 protein expression. This high expression of NLRP3 was also observed in skin fibroblasts by immunofluorescence, in which, NLRP3 was localized in cytosol (Fig. 1B).

**Figure 1.**
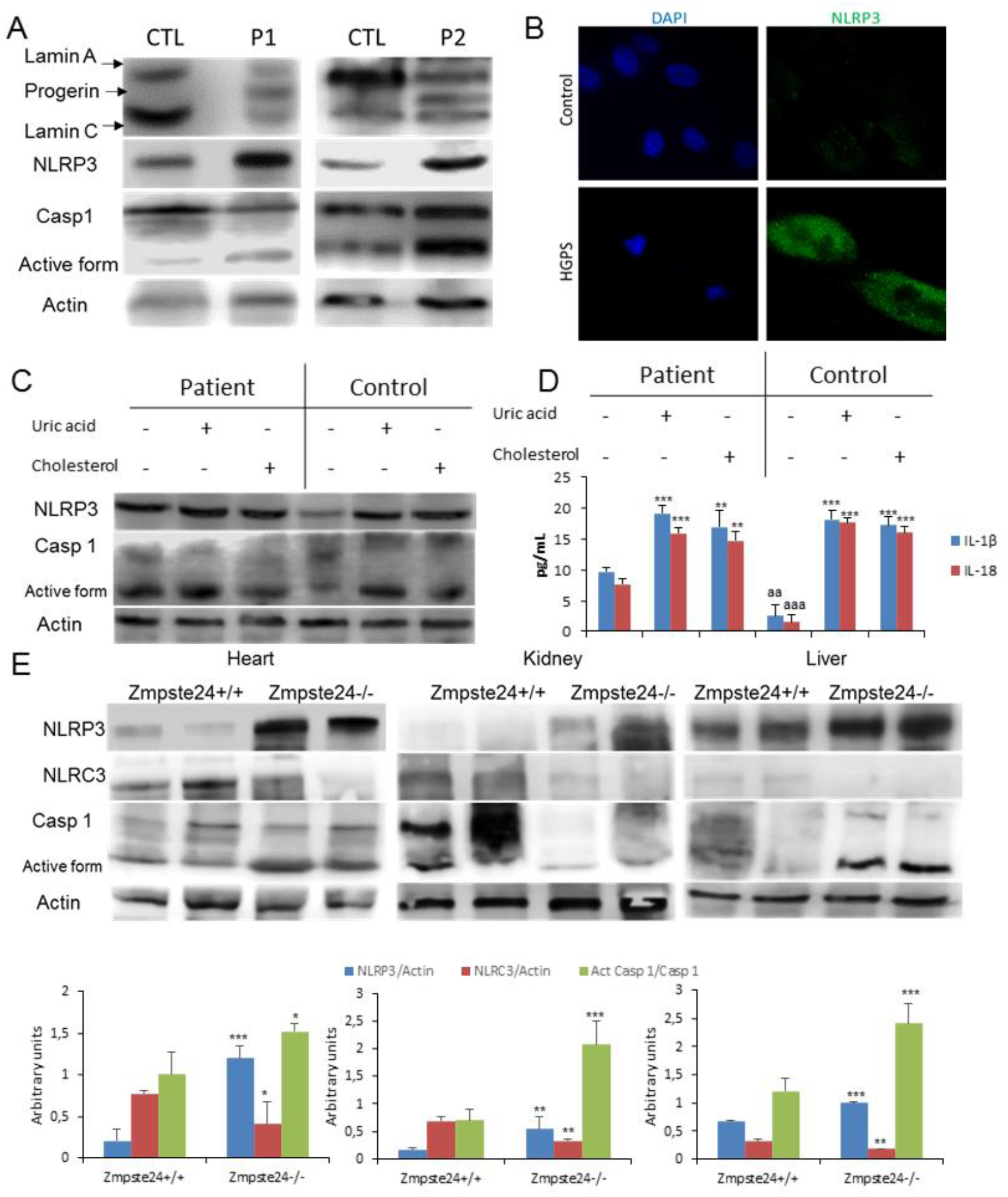
NLRP3 signalling is associated to HGPS. (A) Western blot analysis with representative blot including lamin A/C, NLRP3, caspase 1, and actin levels in skin fibroblats from patients with HGPS, n = 2 controls and 2 patients. (B) Immunofluorescence (IF) visualization of NLRP3 (green) and nuclei (blue) in skin fibroblasts from a representative patient and control. (C) Protein expression of NLRP3 and caspase 1 in lymphocytes from control and one patient after stimulation with uric acid and cholesterol crystal. (D) IL-1β and IL-18 medium release from lymphoblasts. which were assessed after a 24 hr incubation with uric acid and cholesterol. ***P < 0.001, **P < 0.005, *P < 0.05 treatment *vs* no treatment; ^aaa^P < 0.001; ^aa^P < 0.01 control cells *vs* patient cells. (E) Western blot analysis with representative blot including NLRP3, NLRC3, caspase 1, and actin levels in heart, kidney and liver tissues from wild-type and Zmpste24^−/−^ mice. Densitometric analysis is shown as means ± SD, n = 5 mice per group. ***P < 0.001, **P < 0.005, *P < 0.05 wild-type *vs* Zmpste24^−/−^ mice.

It is known that some soluble circulating factors can induce cardiac and metabolic damage and systemic inflammation (12). Furthermore, metabolic alterations have been observed in animal models HGPS with lipid accumulation (13). To evaluate the grade of response of NLRP3-inflammasome complex in HGPS after metabolic alterations, we exposed HGPS lymphoblasts and age- and passage-matched controls to cholesterol crystals and uric acid, two well known metabolic inductor of the NLRP3 activation and associated to aging (1, 4). Interestingly, HGPS cells showed increased basal levels of NLRP3 and caspase 1 protein expression with a moderate increment after cholesterol and uric acid compared to control lymphoblasts (Fig.1C). These high expression was accompanied to increased IL-1β and IL-18 release with high basal levels in HGPS, as well (Fig. 1D).

### Zmpste24-deficient mice show NLRP3-inflammasome hyperactivation

To try to extend these observations to an *in vivo* model, we evaluated the inflammasome complex status in a murine model of progeria. For this purpose, we used a mouse model with the absence of Zmpste24 metalloproteinase which leads to a progeroid phenotype similar to the human premature aging syndrome. After to exam the basal inflammasome levels of diverse tissues from wild-type and Zmpste24^−/−^ mice, we observed an evident increment of the Nlrp3 and caspase 1 protein expression in heart, kidney and liver tissues associated to reduced expression of Nlrc3 (Fig. 1E) which was not observed in other tissues such as muscle and lung (Fig. S1). The specific function of the NLRP3-inflammasome complex in the pathophysiology of HGPS was reinforced by the observation that there were no changes in other inflammasomes (Fig. S2).

### NLRP3-inflammasome inhibition improves progerian phenotype

To examine whether pharmacological inhibition of NLRP3 could be an effective treatment in HGPS, we next assessed the effect of MCC950 on mutant fibroblasts. Control and HGPS fibroblasts from a representative patient were treated at two different doses (0.6 and 1.2mM) of MCC950. The results showed a statistically significant dose-dependent increment of growth rate in patient fibroblasts with cell morphology normalization (Fig. 2A). Western blot analyses also showed that the levels of lamin A and C levels remained relatively constant in both control and HGPS cells treated with MCC950 (to low doses 0.6mM after 48h); conversely, progerin signals in MCC950-treated HGPS cells were decreased accompanied by a reduction of IL-1β and an increment of autophagy protein LC3 (Fig. 2B). Interestingly, the NLRP3 inhibition reduced the frequency of abnormal nuclear morphology in both control and HGPS fibroblasts after 48 hours with a correlative inhibition of NLRP3 expression by immunostaining (Fig. 2C-E).

**Figure 2.**
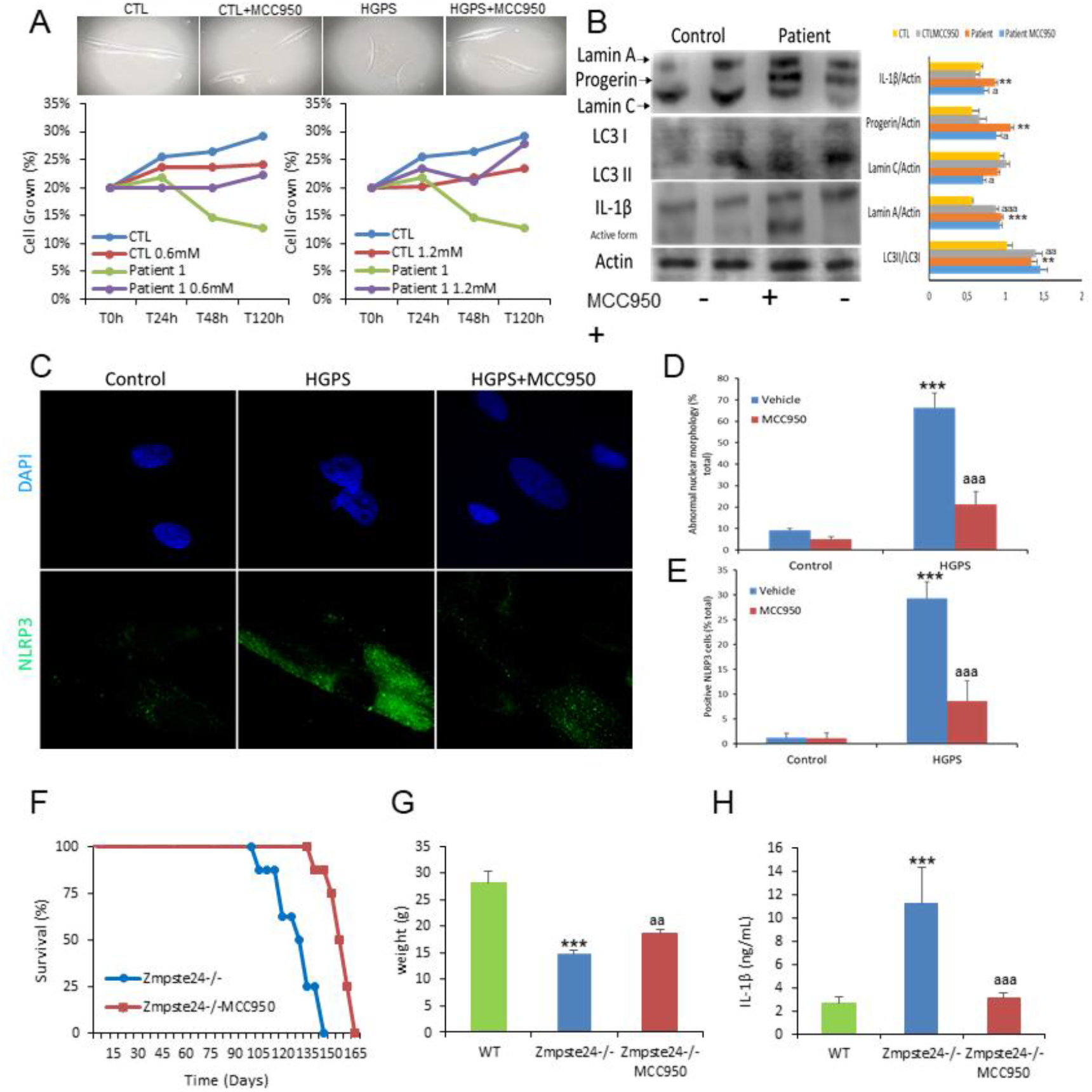
NLRP3 inhibition by MCC950 ameliorates skin fibroblasts and Zmpste24-deficient mice progeroid phenotypes. (A) Cell growth and morphological aspect with MCC950 determined in healthy and representative HGPS fibroblasts. (B) Western blot analysis with representative blots including lamin A/C, NLRP3, caspase 1, LC3 and actin levels in skin fibroblats from control and HGPS patients after 48 hr of vehicle and MCC950 treatment. Densitometric analysis is shown as means ± SD. ***P < 0.001, **P < 0.005, *P < 0.05 no treatment *vs* treatment; ^aaa^P < 0.001; ^aa^P < 0.01 control cells *vs* patient cells. (C and D) Representative fluorescence images of HGPS and control fibroblasts to evaluate the effect of the MCC950 in the nuclear morphology and NLRP3 expression. (E) Kaplan-Meier graph showing a significant increase in the maximum lifespan in WT mice compared with Zmpste24^−/−^ mice. N=7 per group (F) Body weights of the groups over time. (G) Analysis of serum concentrations of IL-1β measured by ELISA. N=6 per group. Data are shown as means ± SD. ***P < 0.001, **P < 0.005, *P < 0.05 wild-type *vs* Zmpste24^−/−^ mice; ^aaa^P < 0.001; ^aa^P < 0.01 vehicle *vs* MCC950.

### NLRP3-inflammasome extend longevity in Zmpste24-deficient mice

Finally, in this work we explored the in vivo effect of the pharmacological inhibition of Nlrp3 in Zmpste24^−/−^ mice. MCC950 was administered by i.p. route at 20mg/kg daily. Interestingly, Zmpste24^−/−^ mice treated with MCC950 significantly extended the longevity of Zmpste24^−/−^ mice with an increment in mean lifespan of 19.2% and in maximum lifespan of 13.9%, improved body weight from 14.7 ± 0.7 g to 18.6 ± 0.7 g (P < 0.001), and reduced IL-1β (Fig. 2F-H).

## Discussion

During the last years, the role of the NLRP3-inflammasome during aging has been subject to study. Genetic deletion of Nlrp3 in mice has been shown to improve lifespan and health by attenuating multiple age-related degenerative changes such as cardiac aging, insulin sensitivity with glycemic control, bone loss, cognitive function and motor performance (4–6). Further, NLRP3 has been studied in cardiovascular diseases. NLRP3 inflammasome is up-regulated in atherosclerosis, myocardial infarction, ischemic heart disease, chronic heart failure, or hypertension (5). In this context, cardiovascular diseases have been shown accelerated in HGPS patients and progerin, the abnormal form of prelamin-A, has also been shown to induces atherosclerosis and cardiac electrophysiological alterations (14). Furthermore, exogenously expressed progerin was showed increases inflammation (15). On the other hand, the AIM2 inflammasome has been shown to be activated after pharmacological alterations of the nuclear envelope integrity compatible with laminopathies (16). For this reasons, it is tempting to speculate that the inflammasomes could contribute to increased aging observed in progeroid syndromes.

During the last years, an effort has been done to find specific pharmacological inhibitors of NLRP3. While several of these inhibitors have been shown to have a specific direct action on NLRP3, others have been shown indirect inhibitory effects (17). MCC950 and analogues are a specific small-molecule inhibitors of the NLRP3, with remarkable therapeutic potential in human diseases.

Many strategies to treat HGPS have been studied but anti-inflammatories have been poorly studied (9). Interestingly, several of these strategies have anti-inflammatory potential and many of these have been shown to induce indirect inhibition of NLRP3 such metformin, resveratrol, rapamycin, quercetin or spermidine (4).

In the current study, we provide evidence showing that the NLRP3-inflammasome complex is an important determinant of HGPS in human cells and mice. We show that the specific expression of NLRP3 and other components of the complex rapidly increase in the skin fibroblasts from patients and the specific tissues associated to the human progeroid phenotype of an animal model such as heart, liver and heart (8,9) where previously were described inflammatory phenotype (10). Interestingly, we also observed reduced levels of Nlrc3 which is a negative regulator of inflammatory signaling pathways and NLRP3 inflammasome (18, 19). Moreover, we show the continuous activation of the NLRP3 inflammasome in immunological cells like lymphocytes from patients. Finally, pharmacological inhibition of NLRP3 by MCC950 treatment improved cell survival and morphology, reduced inflammation and, in HGPS mice resulted in extension of longevity and the prevention of inflammasome-dependent inflammatory event. Therefore, this HGPS treatment strategy, focused on the inhibition of NLRP3-inflammasome complex, could constitute an alternative therapy to slow disease progression in patients with progeria.

## Acknowledgments

This study was supported by a grant from the Andalusian regional government (Grupo de Investigacion Junta de Andalucia CTS113 and Consejería de Salud de la Junta de Andalucia: PI-0036-2014).

## Author Disclosure Statement

The authors declare that no conflict of interest exists for any of them.

---

**Supplementary Figure 1.**
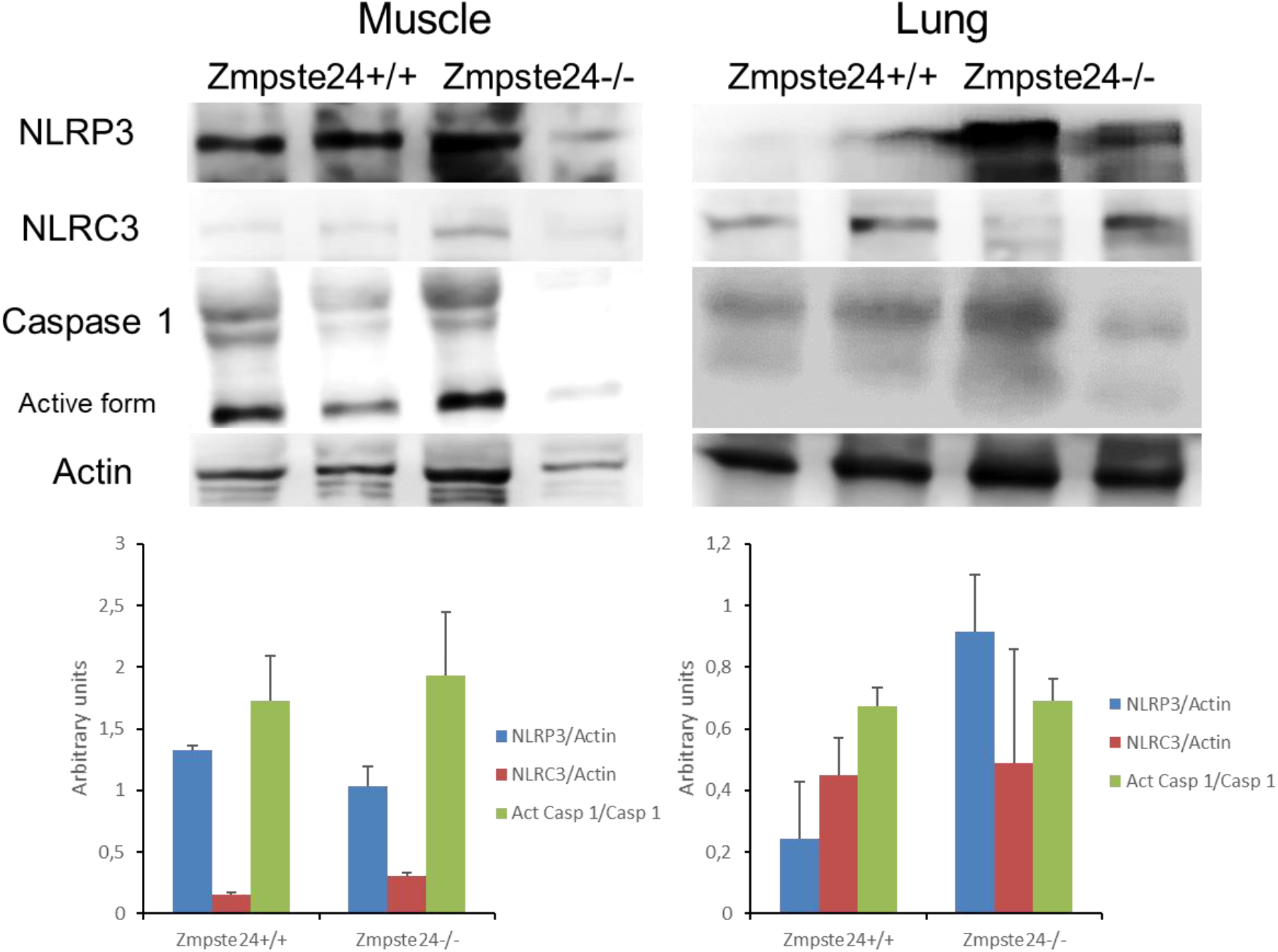
NLRP3-inflammasomes expression in lung and muscle from progeroid animals. Western blot analysis with representative blot including NLRP3, NLRC3, caspase 1, and actin levels in lung and muscle tissues from wild-type and Zmpste24^−/−^ mice. Densitometric analysis is shown as means ± SD, n = 5 mice per group.

**Supplementary Figure 2.**
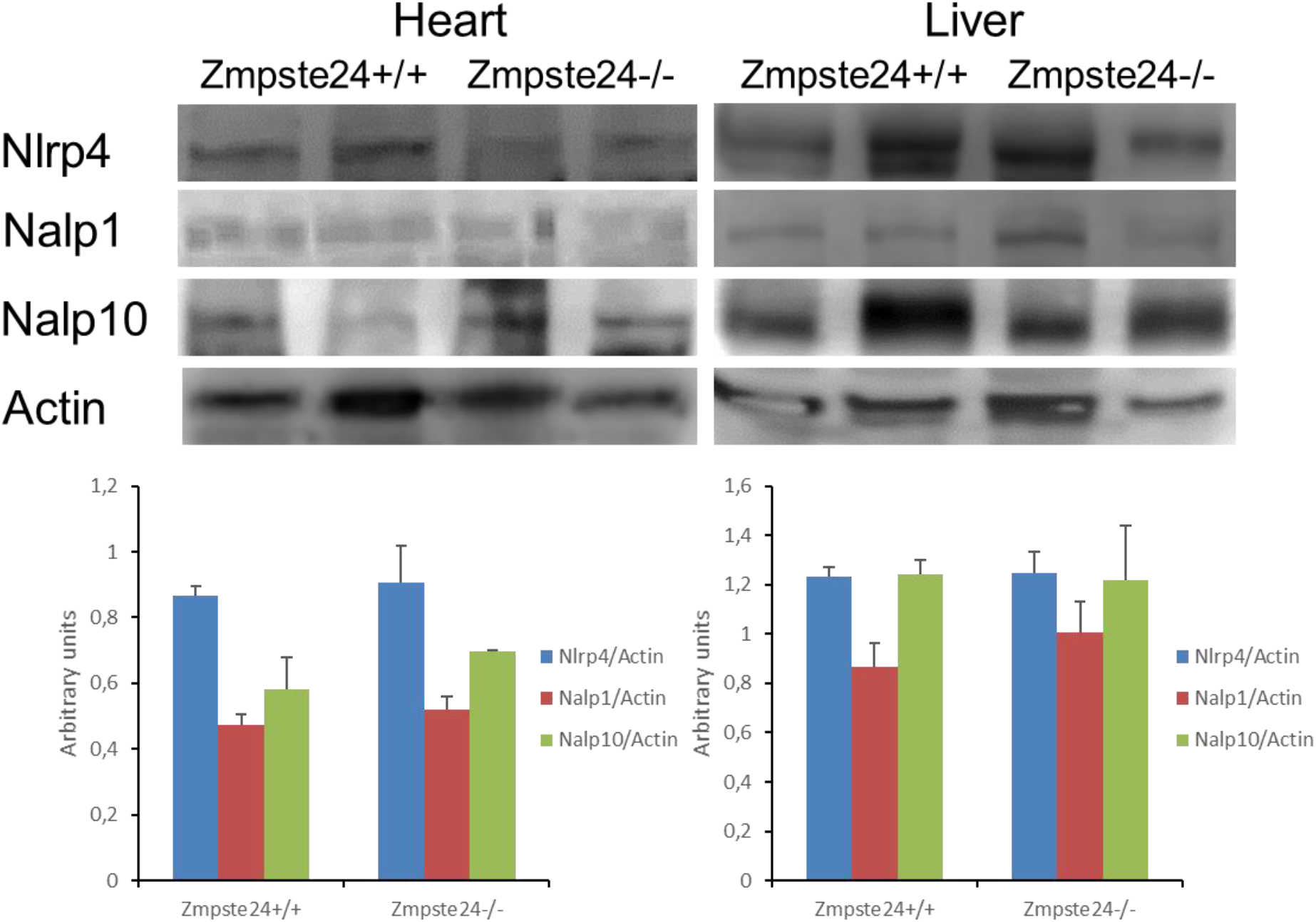
Other inflammasomes are not involved in heart and liver from pathophysiology of progeroid animals. Western blot analysis with representative blot including Nlrc4, Nalp1, Nalp10, and actin levels in heart and liver tissues from wild-type and Zmpste24^−/−^ mice. Densitometric analysis is shown as means ± SD, n = 5 mice per group.

## Notes

### Competing Interest Statement

The authors have declared no competing interest.

